# Effects of Exercise and Konjac on Lipid Metabolism and Mechanism in Obese Rats

**DOI:** 10.1101/2025.03.26.645401

**Authors:** Ling Ruan, Zhengqing Lv, Guanghua Wang, Shoubang Li, Kun You, Menglin Chen, Siyuan Liu, Lu Pu, Xinyue Liu, Yun Pang, Siyu Liu

**Affiliations:** Institute of Physical education, Xi’an Shiyou University, Xi’an, Shaanxi, China; Department of Public Instruction, Xinjiang Vocational University, Urumqi, Xinjiang, China; Xi’an Jingkai Second Middle School, Xi’an, Shaanxi, China

**Keywords:** Konjac, Obesity, Lipid metabolism, Exercise

## Abstract

**Objective:** To study the effects of exercise and konjac on body weight and lipid metabolism in obese rats, and the possible mechanism of exercise combined with konjac on body weight and lipid metabolism.

**Methods:** Fifty male Sprague Dawley rats were randomly divided into 5 groups. After six weeks of intervention, blood lipid, body weight, liver weight, inflammatory factors, oxidative stress markers and proteins were measured.

**Results:** Weight and liver weight in HFD group were significantly higher (P<0.05, P<0.01) than those in CON group. Compared with HFD group, blood lipid index in intervention group were significantly decreased; The levels of MDA (P<0.01, P<0.05, P<0.05) and SOD (P<0.05) in intervention group were significantly decreased; the GSH levels in HAE group and HKE group were significantly increased(P<0.01, P<0.05); the levels of IL-6, TNF-α, and leptin in HAE group (P<0.05, P<0.05, P<0.01) and HKE group (P<0.01, P<0.01, P<0.05) were significantly decreased; ApoA5 in intervention group was significantly increased (P<0.01), SREBP-1c and LXR α was significantly decreased (P<0.01).

**Conclusion:** Konjac or exercise alone can improve the lipid metabolism, and the comprehensive effect is better. The mechanism of regulating lipid metabolism may be related to the expression of LXRα, SREBP-1c and ApoA5.

## Introduction

Obesity occurs due to a dysfunction of lipid metabolism[1].Recently, the prevalence of obesity and associated lipid metabolism disorder has been increased. Obese and lipid accumulation in body are strongly related to inflammory and oxidative stress[2]. Due to the dangers of obesity, people needs nutrition support and exercise training.

Konjac is the common name of Asian plant konjac[3]. Konjac glucomannan (KGM) is a hydrophilic soluble fiber extracted from konjac tubers. It can regulate lipid metabolism by promoting the growth of intestinal mucosa, improving defecation, and changing intestinal microbiota[4]. It has been proved to be a potential auxiliary tool for the treatment of obesity[5].Studies have shown that eating KGM can improve LDL cholesterol concentration, and its potential may be greater than other fibers. In addition, konjac as dietary fiber (DFS) can control body weight by reducing energy intake, reducing energy absorbed from food intake, and increasing postprandial energy consumption (possibly by increasing gastrointestinal peristalsis[6]or increasing excretion of bile acids[7]. This is the same as the results of the controlled intervention study of Clark et al[8]. Epidemiological studies have shown that a higher intake of dietary fibre is associated with smaller body weight and waist circumference[9–10]. This shows that konjac, as a nutritional supplement, has a beneficial effect on weight loss and lipid metabolism in obese people.

Exercise is reported to enhance the ability of skeletal muscles to utilise lipids as opposed to glycogen, thus reducing plasma lipid levels[11]. Lipoprotein lipase activity is also increased by aerobic exercise(AE) which results in clearance of triglycerides(TG) from blood circulation[12].Regular exercise has positive effects on obesity, metabolic dysfunctions, and endothelial dysfunction[13–15]. Exercise are determinant factors for the reversal of metabolic disorders in the experimental model of obese induced by HFD[16].In this study, we hypothesis that combined intervention of konjac and aerobic exercise was improved in rats experimental model of obesity and lipid metabolism induced by HFD. The primary objective of this investigation was to compare the individual and combined effects of konjac and aerobic exercise training on reducing weight gain, and inflammation in obese rats induced by high-fat diet. We hypothesized konjac supplementation would attenuate weight gain like the effects of exercise training, and the combined effects would be better than that of either intervention alone.

## Materials and method

### Institutional Review Board Statement

All the procedures were performed in compliance with the Institute’s guidelines and with the Guide for the Care and Use of Laboratory Animals published by the US National Institutes of Health (NIH Publication No. 85-23, revised 1996). The institutional animal care committee approved the study of GDMLAC. Male 8-week-old Sprague–Dawley rats ranging from 180 to 220 g were obtained from the Guangdong Medical Laboratory Animal Center (GDMLAC) (Guangzhou, China), according to the principles of the Declaration of Helsinki, and the experiments were approved by the Animal Experiment Ethical Inspection Form of Huanan Normal University (IACUU-2008-0020)

### Experimental materials

Experiment design of cohort, body weight (BW) and food intake (FI) were measured every two weeks. After 6 weeks of exercise and konjac feeding, rats were fasted for 12 hours, weighed and anesthetized, blood was drawn from abdominal aorta, and body weight, liver and heart weight were recorded. Part of the liver was placed in 10% formalin for HE staining to observe the pathological changes; Part of the liver was placed in 10% neutral formaldehyde for immunohistochemical test, and part of the liver was placed in liquid nitrogen (-80°C) for Western blot test. Oxidative stress, inflammatory cykocis, and lipid metabolism protein were analysed after 22 weeks(Fig 1).

**Fig 1.**
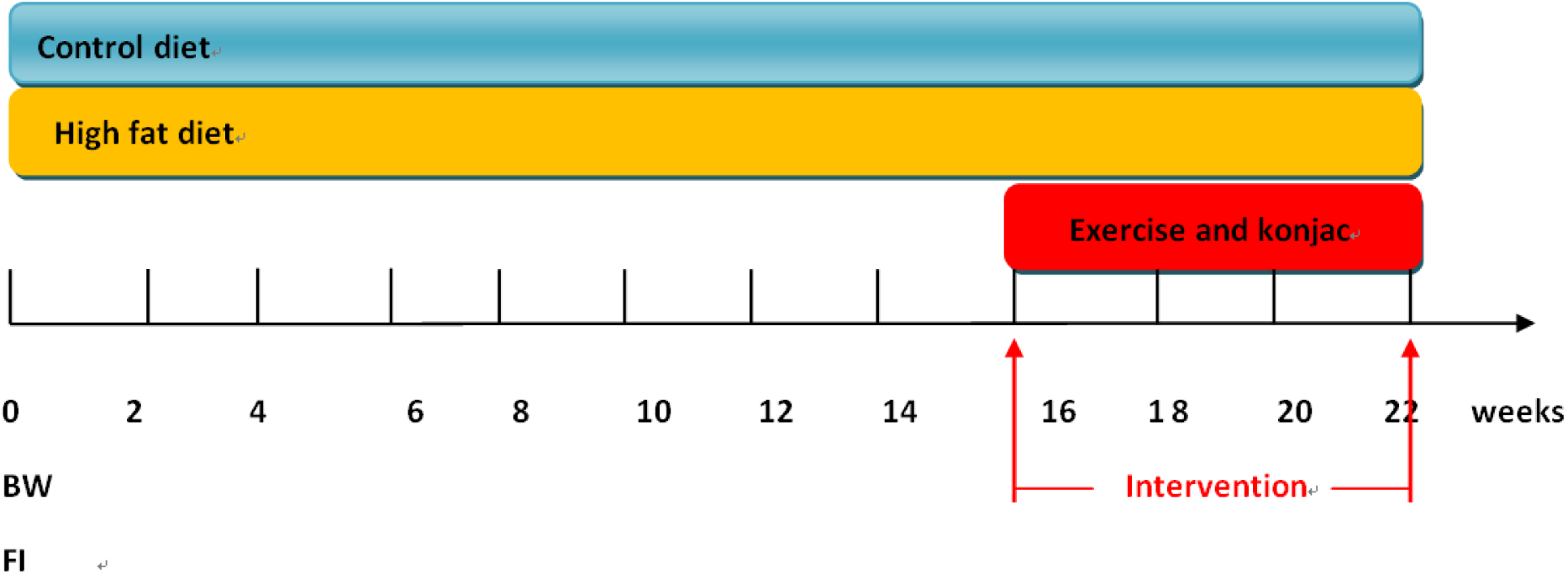
Konjac and/or exercise Intervention treatment.

### Exercise regimen and administration regimen

The setting of exercise load is designed according to the health condition of rats and referring to Bedford TG classic treadmill experimental model[17].Rats in HKM group and HKE group were given KGM at the dose of 50mg/kg body weight every day, the HAE and HKM groups engage in 60 minutes of aerobic exercise per day at an equal speed, 5 times a week. Based on the health status of rats and referring to the Bedford TG classic treadmill experimental model, design exercise load settings. After 6 weeks of exercise and konjac feeding, rats fasted for 12 hours, were weighed and anesthetized, and blood was drawn from the abdominal aorta to record weight, liver, and heart weight. Part of the liver was placed in 10% formalin for HE staining, and pathological changes were observed. Part of the liver was subjected to immunohistochemical testing in 10% neutral formaldehyde, while part of the liver was subjected to Western blot testing in liquid nitrogen (-80°C). Analyze oxidative stress, inflammatory cells, and lipid metabolism proteins using experimental reagents after 16 weeks.

### Animals and treatments

The protocols of this study were approved by the Animal Care and Use Committee of Xi’an Jiaotong University.Male Sprague-Dawley (SD) rats (180-220g, 8 weeks) were purchased from the Experimental Animal Center of Xi’an Jiaotong University (Xi’an, China). All animals were raised in SPF level laboratory animal center (23±1°C, humidity 60-70%, 12 hours light / dark cycle). All rats were fed with free diet, drinking water, providing national standard rodent bedding and standard feed, natural light, ultraviolet light disinfection, and keeping the cage ventilated and dry. After adaptive feeding for 1 week, using systematic random grouping method, all rats were numbered alternately and randomly divided into experimental group and control group. SD rats (n=50) were randomly divided into two groups: normal control group (CON; n=10); High fat diet group (HFD; n=40) after adaptive feeding for 1 week. The high-fat diet group rats were fed a high-fat diet with 5% sucrose, 18% lard, 15% egg yolk powder, 0.5% sodium cholate, and 1% cholesterol added to the basic diet. Rats were fed with high-fat diet, at the end of 16 week, the rats whose body weight was more than 10% of the normal group were regarded as successful rats model[18].After successful modelling, 40 rats fed with high-fat diet were randomly divided into 4 groups: high-fat diet group (HFD; n=10), aerobic exercise group (HAE; n=10), KMG group (HKM; n=10), and aerobic exercise combined with KMG group (HKE; n=10). Among them, 47 rats were successfully established, 3 failed to reach the standard, and the model forming rate was 94%, and food-grade KGM (purity 90%) was purchased from Shaanxi Mingfu Biotechnology Co. Ltd. in China. The reagent KE was provided by the Drug Inspection Institute of the Nanjing Military Region, with a konjac glucomannan content of 82.4%. Insulin was bought from Sigma Company, USA; Lipid measured reagent kits (Dongou Biology Project Company, China), Insulin measured reagent kits (China Institute of Atomic Energy). The enzyme-linked immunosorbent assay (ELISA) kits for PPARα, ApoA5 and SREBP-1c were obtained from Cusabio Biotech Co. Ltd. (Wuhan, China).

### Serum sample preparation and detection

Rats were fasted for 12 hours and weighed, anesthetized with chloral hydrate (Chengdu Kelong Chemical Reagent Factory, China) by intraperitoneal injection. Then blood samples were collected from the femoral artery and placed in centrifuge tubes without anticoagulant for 30–60 min at room temperature. The serum was then separated by centrifugation at 4000 rpm for 15 min and was stored in a refrigerator at -20°C for further serum index measures. The levels of serum TC, TG, LDL-c, HDL-c, ALT and AST were measured according to the manufacturer’s instructions using microplate reader (BioTek Instruments, Inc., USA). The content of inflammatory factors was detected by ELISA.

### Tissue preparation and index detection

All rats were quickly dissected after blood collection. The liver was washed with cold saline solution and weighed with an analytical balance (Beijing Sartorius Instrument System Co, Ltd.). Then, a small part of the liver was fixed with 10% formalin solution to observe the pathological changes by hematoxylin-eosin (HE) staining, whereas the other parts were stored in a refrigerator at -80°C. The levels of malondialdehyde (MDA) and glutathione peroxidase (GSH PX) in some liver tissues were determined by colorimetry; superoxide dismutase (SOD) in liver tissues was determined by WST-1 method; catalase (CAT) in liver tissues was determined by visible light method. The content of inflammatory factors was detected by ELISA kit, and the standard operation was carried out in strict accordance with the instructions of the kit. Western blot was used to detect the expression levels of PPARα, APOA5, SREBP-1c and LXRα. All operations were carried out in accordance with the manufacturer’s instructions for using the micro board reader (BioTek instruments, USA).

### Statistical analysis

SPSS 21.0 was used to analyze the data. Graphpad prism (version 7.0) was used to make a graph, and the mean±standard deviation (M±SD) was used. All the indexes were analyzed by Two-way ANOVA. The values of P<0.05 or P<0.01 respectively indicated significant differences and significant differences. The values of P<0.05 or P<0.01 respectively indicated significant differences and significant differences.

## Result

### The Change of Body weight, organ weight in each group

As shown in Fig 2 and Fig 3, the initial weight in HAE group was decreased (P<0.05) compare to HFD group, the initial weight in HKE group was decreased (P<0.05) compare to HKM group. And heart weight in HKE group was decreased compare to CON (P<0.05), HKM (P<0.05) group. Compared with CON group, the final weight of HFD group increased (P<0.05), while that of HKE group decreased (P<0.05), compared with HFD group, HKM group, HAE group and HKE group all had lower final body weight(P<0.01, P<0.05, P<0.01).Compared with CON group, the liver weight of HFD group and HKM group increased significantly (P<0.01), compared with HFD group, the liver weight of HKE group was decreased(P<0.05).As shown in Figure3B, there was no significant difference in food intake at each group

**Fig 2.**
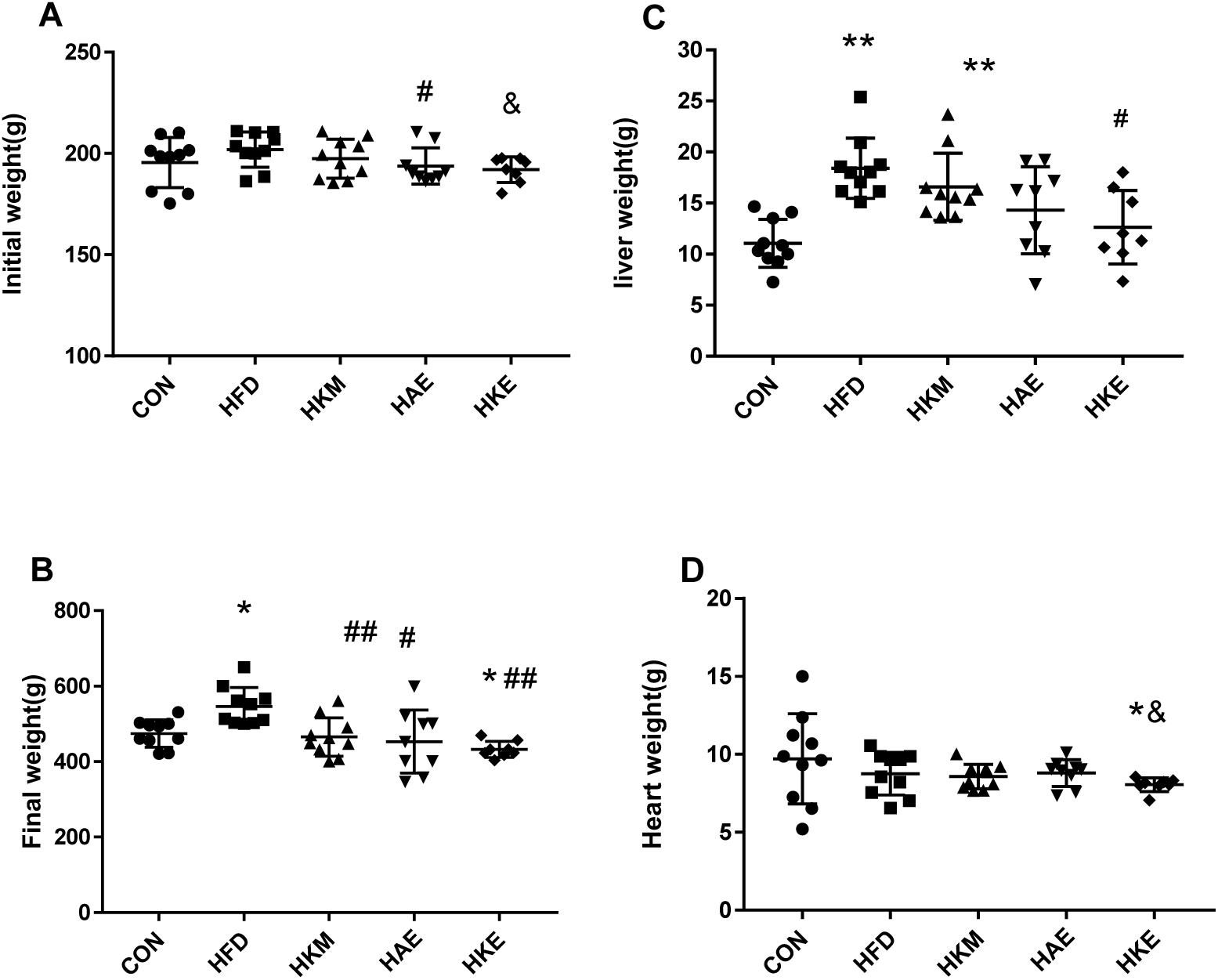
Initial weight(A), Final weight(B), Liver weight (C), and Heart Weight (D) in each group. Note: Value are reported as the means± SD,\**P*<0.05, \*\**P*<0.01 was compare to CON group; # *P*<0.05, ## *P*<0.01 was compare to HFD group; & *P*<0.05, &&*P*<0.01 was compare to HKM group; % *P*<0.05, %% *P*<0.01 was compare to HAE group.

**Fig 3.**
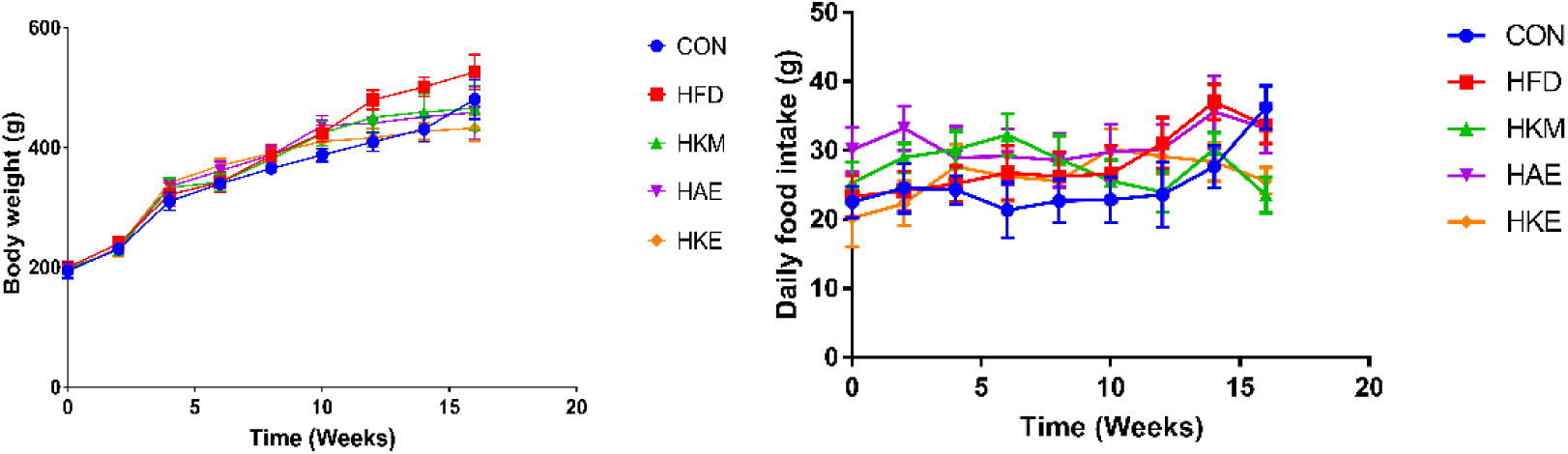
Body weight and daily food intake. Values represent the means±SD. Measurement of body weight in CON, HFD, HKM, HAE, and HKE group rats and dairy food intake. Body weight and food intake from the 0th to 16th week evaluated using two-way ANOVA with konjac and exercise as factors, follow by Bonferroni’s post-test when appropriate.

### Hematoxylin-eosin staining of perirenal adipose tissue

Figure 4 shows the changes of HE-stained perirenaladipose tissue of rats in different groups under light microscope. The size and distribution of adipocytes in the CON group were uniform, and the adipocytes in the HFD group were significantly enlarged, arranged irregularly, and had inflammatory infiltration. After the intervention of exercises and konjac, the volume of fat cells decreases the degree of inflammation infiltration is significantly reduced in all the intervention groups, and the effect of exercise and konjac intervention is more significant.

**Fig 4.**
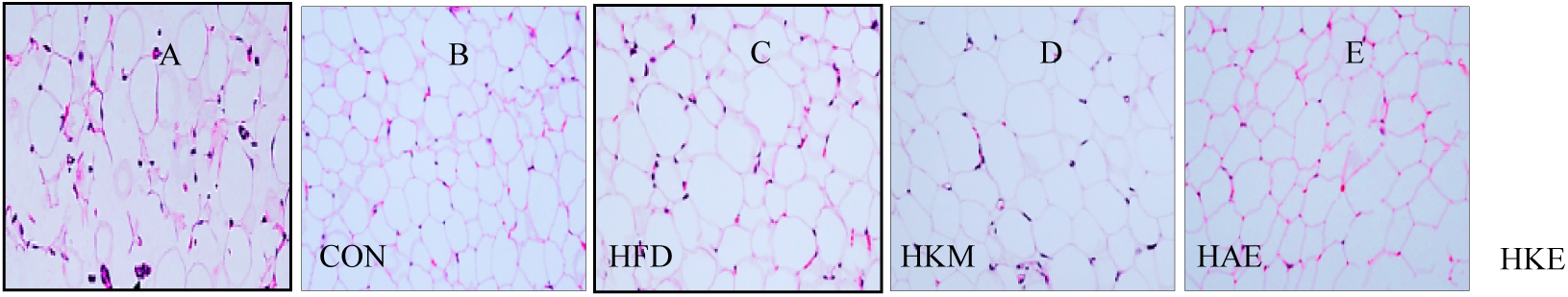
The Optical Microscope Image of HE Dyed in Rat Adipose cells (×400) (A-E)

### The Change of liver lipids, liver enzyme and insulin sensitive in each Group

Table 1 shows that compared with CON group, TG in HFD group was significantly increased (P<0.01), and that in HAE group was increased (P<0.05), compared with HFD group, TG of HKM group was decreased (P<0.05), and that of HKE group was significantly decreased (P<0.01), compared with HAE group, TG in HKE group decreased(P<0.05). Compared with CON group, TC in HFD group, HKM group and HAE group was increased(P<0.05). Compared with CON group, LDL-c in HFD group, HKM group, HAE group and HKE group was significantly increased (P<0.01), compared with HFD group, LDL-c in HKM group, HAE group and HKE group was significantly decreased(P<0.01), compared with HKM group, the LDL-c of HAE group and HKE group was significantly decreased(P<0.01), compared with HAE group, the LDL-c of HKE group was decreased(P<0.05).Compared with CON group, HDL-c in HFD group and HKM group decreased significantly(P<0.01, P<0.05). Compared with CON group, FFA in HFD group, HKM group, HAE group and HKE group was significantly increased (P<0.01), compared with HFD group and HAE group, FFA of HKE group was decreased (P<0.05). Compared with CON group, ALT in HFD group, HKM group and HAE group was significantly increased(P<0.01), compared with HFD group, ALT in HAE group and HKE group decreased significantly(P<0.01, P<0.05),compared with HKM group and HAE group, ALT in HKE group decreased(P<0.05).Compared with CON group, HFD group and HAE group, AST in HKE group was increased (P<0.01, P<0.05, P<0.05). Compared with CON group, fast blood glucose in HFD group, HKM group and HAE group was significantly increased(P<0.01),compared with HFD group, the fast blood glucose in HAE group and HKE group was significantly decreased(P<0.01),compared with HKM group and HAE group, the fast blood glucose in HKE group was significantly decreased(P<0.01, P<0.05). Compared with CON group, fast serum insulin in HFD group was significantly increased(P<0.01), compared with HFD group, fast serum insulin in HKM group, HAE group and HKE group was significantly decreased(P<0.01, P<0.05, P<0.01). Compared with CON group, HOMA-IR of HFD group, HKM group and HAE group was significantly increased (P<0.01), compared with HFD group, HOMA-IR of HKM group, HAE group and HKE group was significantly decreased (P<0.01), compared with HKM group and HAE group, HOMA-IR of HKE group was decreased (P<0.05).

**Table 1.**
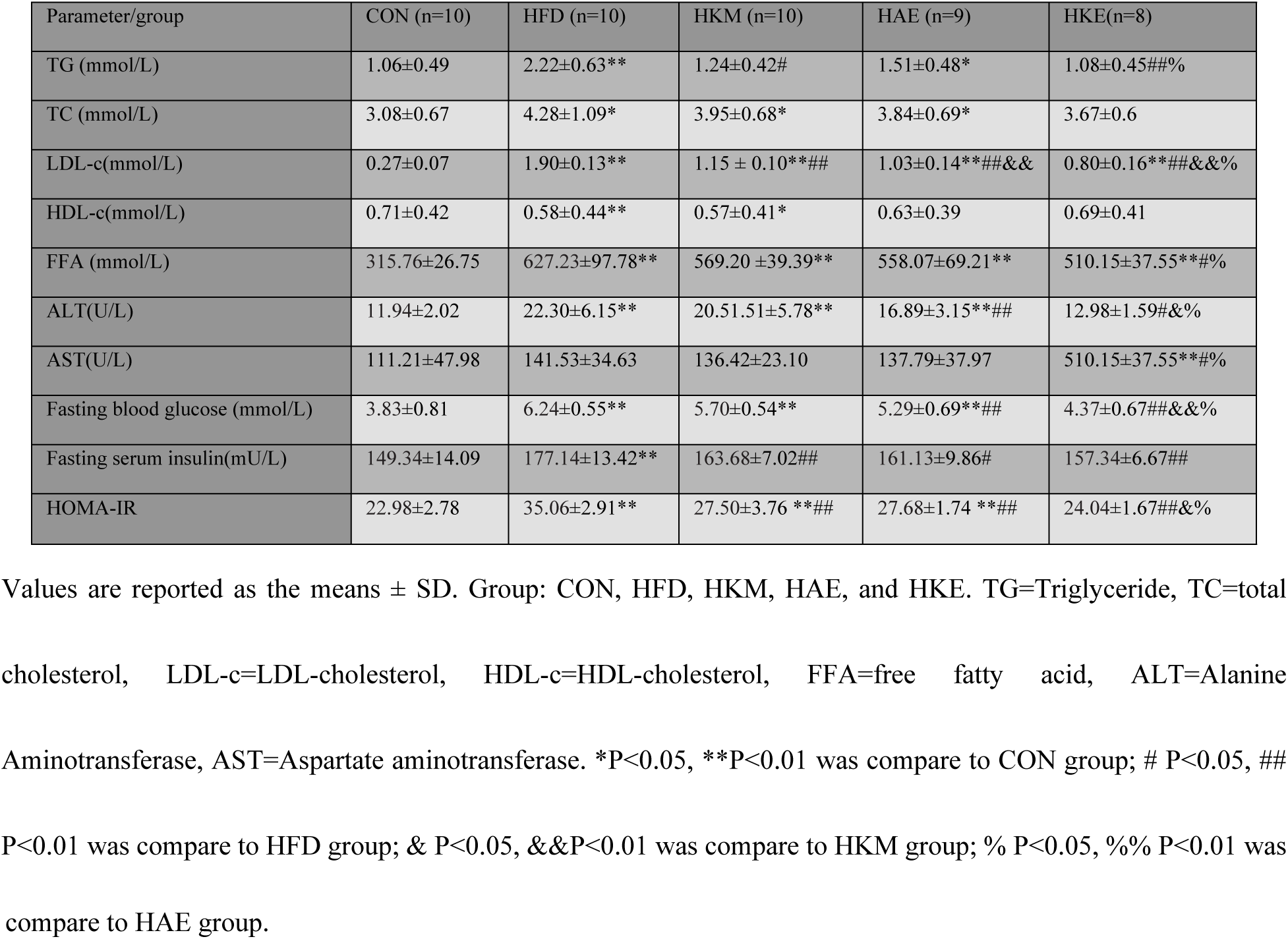
The Change of liver lipids, liver enzyme and insulin sensitive in each Group.

### Oxidative Stress

As Fig 5 show that compared with HFD group, SOD in HKM group increased(P<0.05). Compared with CON group, MDA in HFD group was significantly increased(P<0.01),compared with HFD group, MDA in HKM group, HAE group and HKE group was significantly decreased(P<0.01, P<0.05, P<0.05). Compared with CON group, CAT in HFD group was significantly decreased(P<0.01),compared with HFD group, CAT in HKM group, HAE group and HKE group was significantly increased(P<0.01).Compared with CON group, the GSH of HFD group was decreased(P<0.05),compared with HFD group, GSH in HAE group and HKE group was significantly increased(P<0.01, P<0.05).

**Fig 5.**
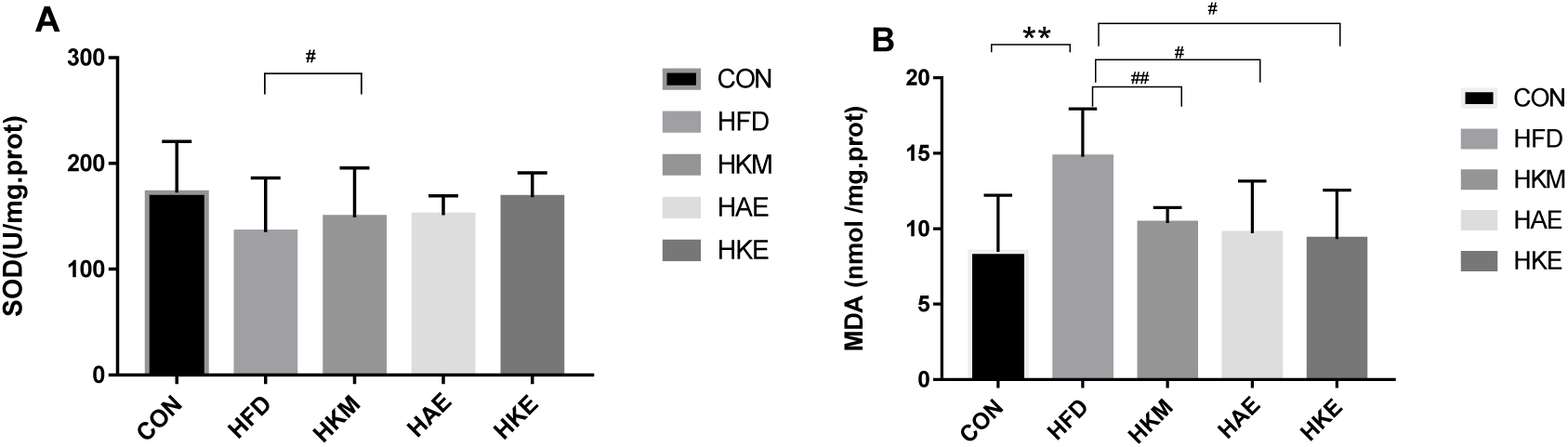
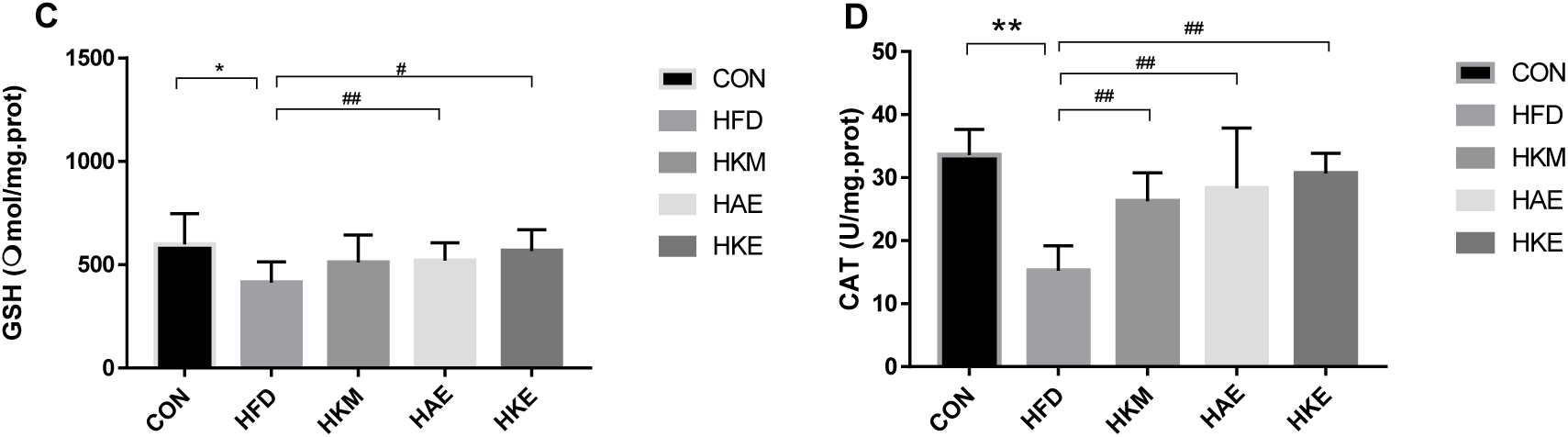
Antioxidant index. Note: Values are reported as the means ± SD.\**P*<0.05, \*\**P*<0.01 was compare to CON group; # *P*<0.05, ## *P*<0.01 was compare to HFD group; & *P*<0.05, &&*P*<0.01 was compare to HKM group; % *P*<0.05, %% *P*<0.01 was compare to HAE group.

### The change of inflammatory factors in each group

The results in Fig 6 show that compared with CON group, the levels of IL-6 in HKM and HKE groups were increased (P<0.05), compared with HFD group, the level of IL-6 in HAE group and HKE group was significantly decreased (P<0.05, P<0.01). Compared with CON group, the release of TNF-α in HFD group and HKM group was significantly increased(P<0.01),compared with HFD group, TNF-α release in HAE group and HKE group was significantly decreased(P<0.05, P<0.01),compared with HKM group, the release of TNF-α in HKE group was significantly decreased(P<0.01).Compared with CON group, leptin release was significantly increased in HFD group, HKM group, HAE group and HKE group(P<0.01, P<0.01, P<0.05, P<0.01),compared with HFD group, leptin release in HAE group and HKE group was significantly decreased(P<0.01,P<0.05),compared with HKM group, leptin release in HAE group was decreased(P<0.05).Compared with CON group, the release level of CRP in HFD group and HKM group was significantly increased (P<0.01, P<0.01).

**Fig 6.**
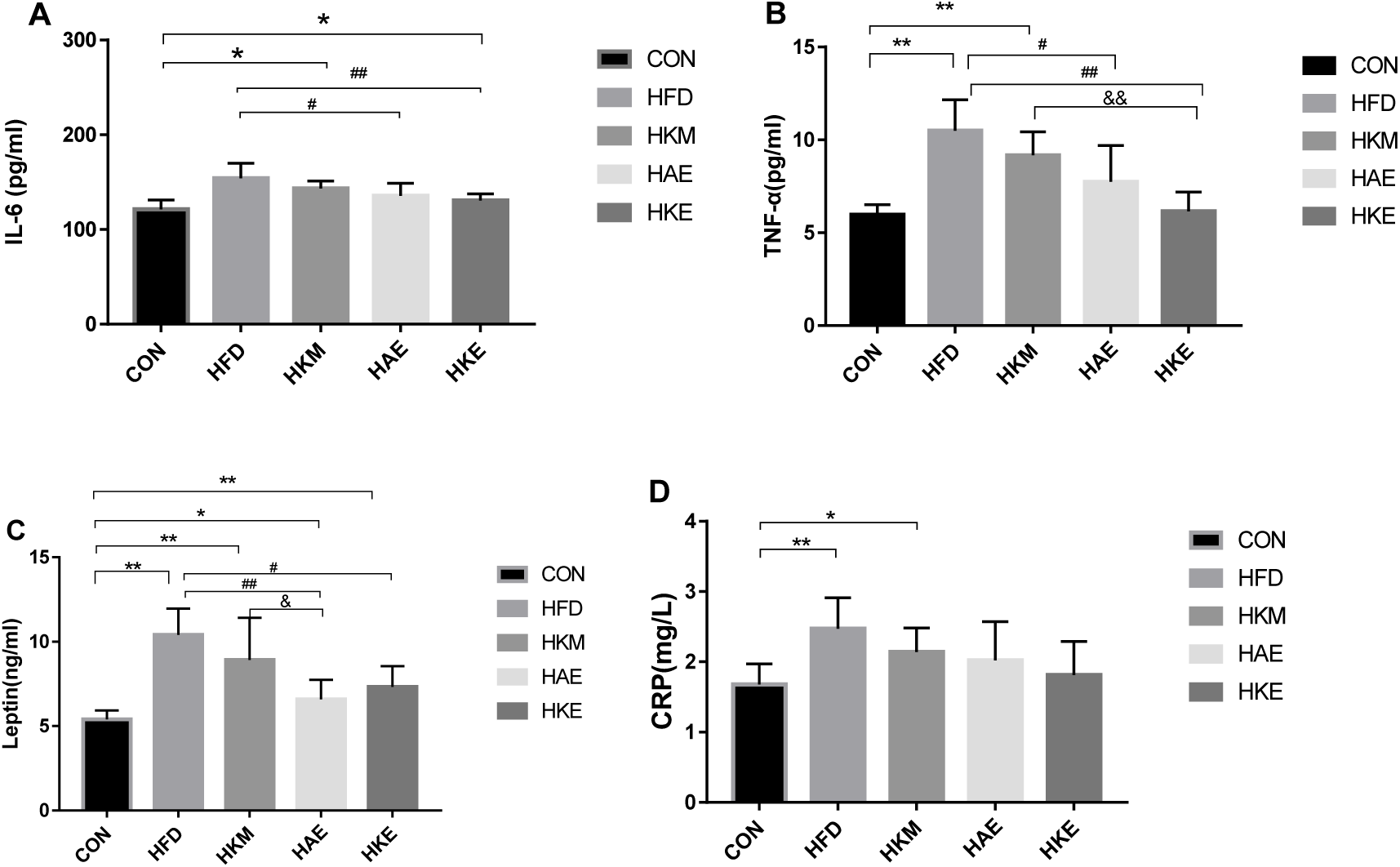
The Change of Inflammatory factors in Each Group. Note: Values are reported as the means ± SD.\**P*<0.05, \*\**P*<0.01 was compare to CON group; # *P*<0.05, ## *P*<0.01 was compare to HFD group; & *P*<0.05, &&*P*<0.01 was compare to HKM group; % *P*<0.05, %% *P*<0.01 was compare to HAE group.

### ApoA 5, PPARα, SREBP-1c, and LXRα contents in the liver tissue

The results in Figure7 show that compared with CON group, the expression of ApoA5 in HFD group, HKM group, HAE group and HKE group was significantly decreased(P<0.01), compared with HFD group, the expression of ApoA5 in HKM group, HAE group and HKE group was significantly increased(P<0.01),compared with HKM group and HAE group, the expression of ApoA5 in HKE group was significantly increased(P<0.01).Compared with CON group, the expression of PPARα in HFD group was significantly decreased(P<0.01),compared with HKM group, the expression of PPARα in HKE group was increased(P<0.05).Compared with CON group, SREBP-1c expression in HFD group, HKM group, HAE group and HKE group was significantly increased(P<0.01), compared with HFD group.SREBP-1cexpression in HKM group, HAE group and HKE group was significantly decreased(P<0.01),compared with HKM group, SREBP-1c expression in HAE group and HKE group was significantly decreased (P<0.05,P<0.01). Compared with CON group, the expression of LXRα in HFD group was significantly increased (P<0.01), compared with HFD group, the expression of LXRα in HKM group, HAE group and HKE group was significantly decreased (P<0.01).

**Fig 7.**
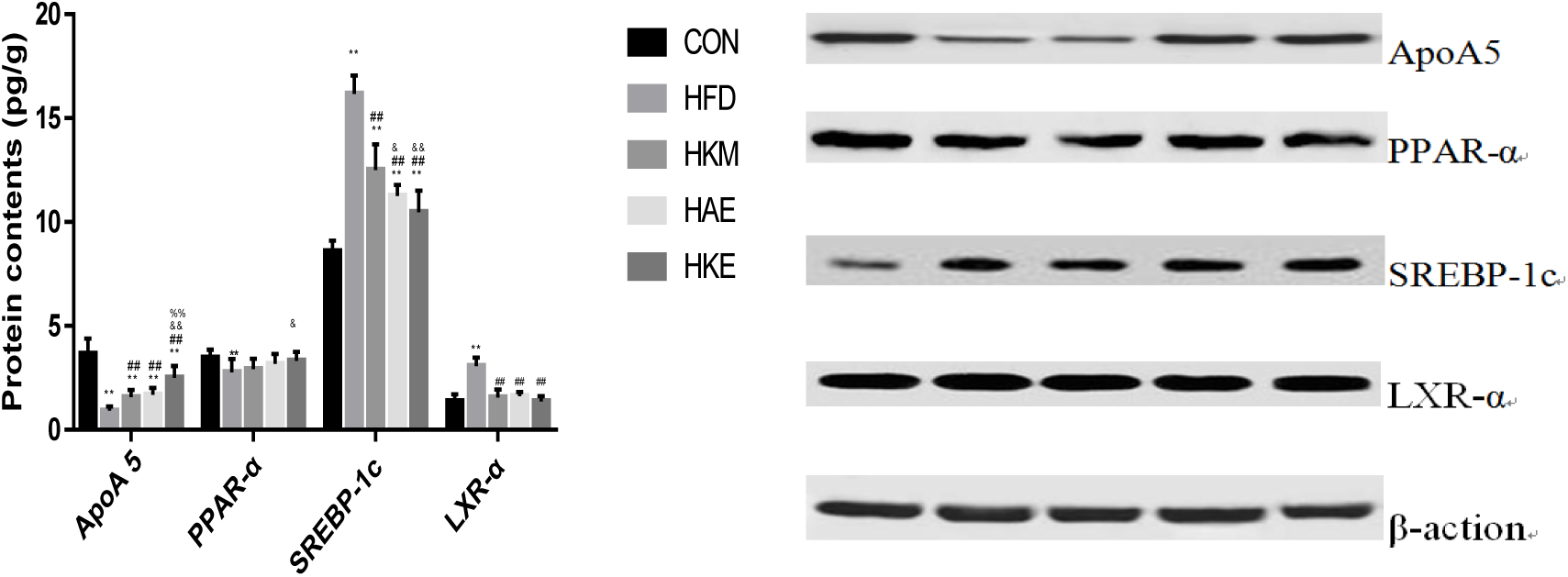
Lipid metabolism protein. Note: Values are reported as the means ± SD.\**P*<0.05, \*\**P*<0.01 was compare to CON group; # *P*<0.05, ## *P*<0.01 was compare to HFD group; & *P*<0.05, &&*P*<0.01 was compare to HKM group; % *P*<0.05, %% *P*<0.01 was compare to HAE group.

### Correlation analysis of ApoA 5, PPAR α, SREBP-1c, LXR α protein and blood lipid

As shown in Fig 8, ApoA5 and PPARα were negatively correlated with TG(P<0.01, P>0.05), TC(P<0.01, P>0.05), FFA(P<0.01, P<0.01)and LDL-c (P<0.01, P<0.01), and positively correlated with SREBP-1c(P<0.01, P<0.01), LXRα(P<0.01, P<0.01)and HDL-c(P>0.05, P>0.05), ApoA5 was negatively correlated with PPARα (P<0.01); SREBP-1c and LXRα were positively correlated with TG(P<0.01, P<0.01), TC(P<0.01, P>0.05), FFA(P<0.01, P<0.01), LDL-c(P<0.01, P<0.01) and negatively correlated with HDL-c(P>0.05, P>0.05),SREBP-1c was negatively correlated with LXRα(P<0.01); TG and TC were negatively correlated with FFA(P<0.01, P<0.01), LDL-c(P<0.01, P<0.01), positively correlated with HDL-c (P>0.05, P<0.01), TG was negatively correlated with TC(P<0.05); FFA was positively correlated with LDL-c(P<0.01) and positively correlated with HDL-c(P>0.05); There was a positive correlation between LDL-c and HDL-c (P>0.05).

**Fig 8.**
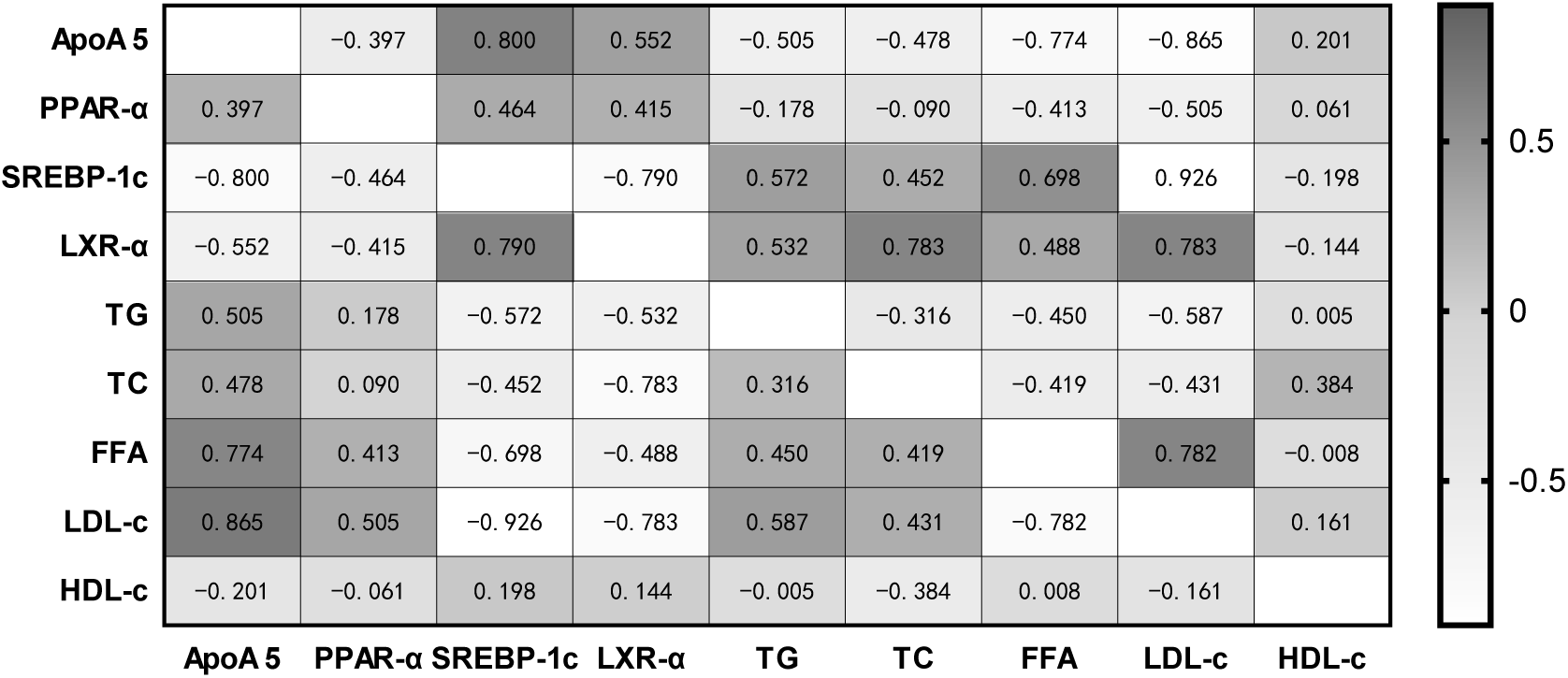
Heatmap Analysis of Pearson Correlation between the Protein Expression of ApoA5, PPARα, SREBP-1c, LXRα and blood lipid.

## Discussion

There is strong evidence that regular exercise helps to lose weight and fat, so as to improve the metabolic health of obese patients. Gasier et al. have found that exercise can effectively inhibit the bodyweight growth of obese rats induced by a high-fat diet, and improve lipid metabolism disorder[19].Meanwhile, konjac as a natural polymer soluble dietary fiber, has been proved to play an important role in defecation, weight loss, blood glucose and blood lipid[20].This may be because konjac can reduce the gastric emptying rate of food, delay the digestion and absorption of nutrients, especially the absorption of monosaccharides, inhibit the synthesis of fatty acids in the body, enhance gastrointestinal peristalsis, increase defecation volume, and reduce the reabsorption of nutrients and water in food residues by the gastrointestinal tract[21].Our results showed that compared with HFD group, the weight of rats in HAE and HKM group decreased significantly; At the same time, our results also indicate that after 16 weeks of high-fat diet, the fasting serum insulin, leptin and HOMA-IR levels in HFD group are higher than those in CON group. High fat diet can increase blood glucose, stimulate insulin secretion and insulin resistance, promote liver synthesis of TG and LDL-C, and increase plasma TG and LDL-C concentrations[22], but after exercise and konjac intervention, the results of these test indicators were lower than those of the HFD group. This shows that exercise and konjac can not only reduce the weight of rats, but also affect their lipid metabolism, especially the levels of TG and LDL-C (P<0.01), regulate leptin and improve insulin sensitivity.

High fat diet causes oxidative stress in rats[23]. Oxidative stress refers to a state of imbalance between oxidation and antioxidation in vivo. Stimulation of high FFA and LDL-c can promote the production of ROS. Exercise can regulate the oxidative state of the body by reducing weight gain and fat accumulation[24]. The results showed that compared with the HFD group, the HAE and HKE groups had higher levels of SOD, CAT, and GSH, and lower levels of MDA. This shows that aerobic exercise reduce body weight and fat accumulation, and regulate the oxidation state by increasing SOD, CAT, GSH and significantly reducing MDA.At the same time, the levels of ALT and AST in HAE group decreased significantly, indicating that exercise can reduce the damage of oxidative stress caused by lipid deposition to hepatocytes, inhibit the occurrence and development of IR, increase insulin sensitivity and improve blood glucose. Studies have shown that konjac also has a certain antioxidant effect[25]. KGM can reduce the production of peroxide, normalize the activities of peroxidase and oxidized glutathione reductase, restore GSH level and SOD activity, so as to enhance the function of free radical antioxidant defence system, which is stronger than the classical antioxidants vitamin E and vitamin C.KGM reduces MDA mainly by increasing the levels of antioxidant SOD, GSH and CAT in liver, thus reducing the production of free radicals, which is consistent with our results.

Due to the metabolic disorder of adipocytes, the immune response in vivo is abnormally activated, which leads to the occurrence of inflammatory reaction, and the level of many pro-inflammatory factors will also increase. Under the condition of obesity, inflammatory factors such as TNF-α and IL-6 can phosphorylate I–κB and dissociate it from NF-κB, thus activating NF-κB. Activates and regulates the gene transcription of various inflammatory factors and related inflammatory substances, further promotes the synthesis and secretion of TNF-α and IL-6, aggravates the inflammatory reaction, expands the inflammatory reaction, and finally leads to insulin resistance. The over expression of TNF-α and IL-6, which was related to many inflammatory diseases was an important hub connecting inflammation and insulin resistance[26–27].In general, inhibition of TNF-α and IL-6 secretion is considered to be an effective strategy for controlling inflammatory diseases[28].CRP is a sensitive indicator of low-level inflammatory response, and high-fat diet can lead to increased CRP level and induce adipose tissue inflammation in rats[29].Different studies have shown that both exercise and konjac have anti-inflammatory effects[30–31]. Exercise can inhibit the expression of inflammatory factors by increasing the level of glucocorticoid during exercise; or reduce the secretion of inflammatory factors TNF-α and IL-6 in endothelial cells by improving endothelial function; and reduce the synthesis of TNF-α and IL-6 in adipose tissue due to atrophy of adipose tissue, decrease of body fat and decrease of BMI after exercise[32]. KGM in konjac also improves insulin resistance by lowering levels of inflammatory factors. In our experiment, TNF-α and IL-6 were significantly increased in the HFD group compared with the CON group, while TNF-α and IL-6 were significantly lower in the HAE and HKM groups than in the HFD group, suggesting that exercise and konjac can effectively reduce inflammation induced by high fat diet.

Through our study and comparison, we found that the level of final weight and liver weight in the HKE group decreased significantly compared to the HFD group; it’s much lower than the HAE group and HKM group. Our results show that the combination of konjac and exercise has a better effect on obese rats than single intervention. This better effect also includes effectively reducing the blood lipid level, antioxidant effect, anti-inflammatory effect and improving insulin resistance in rats. In conclusion; our data seem to indicate that the combination of exercise and konjac has a better effect on lipid metabolism in rats.

To further explore the underlying molecular mechanisms, the protein expression of some important genes involved in metabolism (protein) and inflammatory factors was detected in our study.

PPARα is transcription factors that modulate the expression of genes involved in lipid metabolism, energy homeostasis and inflammation[33].Experimental evidence shows that PPARs is the master regulator of hepatic[34]. High fat diet induced obese are partly due to up-regulation of the PPARα mediated metabolic pathways. In addition to activation of PPARα, partial activation of PPARγ also has beneficial effects, mainly mediated by increased adiponectin expression and decreased insulin resistance[35].Notably, our results suggest that exercise and konjac induce PPARα and PPARγ, which may be potential target genes for exercise and konjac mediated moderating the progression of obesity.

LXRα plays an important regulatory role in liver lipid metabolism by participating in the reverse transport of cholesterol, promoting the conversion of cholesterol into bile acid in the liver with bile excretion, inhibiting the synthesis of cholesterol in the liver, and regulating the synthesis of fatty acids and phospholipids in the liver[36].Kazeminaab et al. found that 4-week aerobic exercise can reduce the expression of the LXRα gene and increase HDL-c, which leads to the depletion of intracellular cholesterol. Therefore, exercise can reduce the content of cholesterol by regulating the expression of LXRα[37].Our study showed that LXRα was positively correlated with TC, while LXRα was negatively correlated with HDL-c (P< 0.01), which is consist with his result. The increased expression of LXRα has also been shown to be related to the degree of inflammation. Webb et al. reported that LXRα may be involved in redox-sensitive signal transduction and reduce oxidative damage[38], while activation of LXRα mediated by increased insulin sensitivity and insulin secretion leads to decreased blood glucose levels[39].

ApoA5 can regulate TG metabolism through various mechanisms, one of which is to increase the activity of Lipoprotein Lipase (LPL) and Hepaticlipase (HL) to promote TG hydrolysis[35,40].The gene and protein expression of ApoA5 is regulated by a variety of nuclear receptor and non-nuclear receptor factors. Among these factors, PPARα and LXRα are two key nuclear receptors that can up-regulate and down-regulated, respectively, the expression of ApoA5.

It has been found that over expression of SREBP-1c can activate transcription of targeted fatty acid synthase (FAS) gene, induce increased lipid production and fatty acid content, and lead to lipid accumulation in non-adipose tissues such as liver, resulting in glucose and lipid metabolism disorders and obesity[41].After hepatic insulin resistance, the LXRα/RXR signaling pathway was upregulated, and SREBP-1c gene transcription was increased. LXRα can directly or indirectly regulate the synthesis of fatty acids and TG in liver by lipogenic genes such as SREBP-1c, and regulate phospholipid and lipoprotein by activating phospholipid transporter protein (PLTP) and lysophosphatidylcholine acyltransferase 3 (LPCAT3), resulting in the increase of new fat in liver. The over expression of LXRα can cause the accumulation of new fat in the liver leading to hyperlipidemia, nonalcoholic fatty liver disease and other diseases, which is harmful to human health[42]. LXRα is mainly involved in the process of liver lipid production. Our data showed that SREBP-1c (P<0.01) and LXRα (P<0.01) levels were significantly lower than those in the HFD group after konjac and exercise intervention. However, ApoA5 (P<0.01) and PPARα levels increased significantly. So, we speculate that the mechanism of lipid metabolism regulation by exercise and konjac may decrease the LXRα and SREBP-1c protein expression levels to increase the ApoA5 expression and then metabolize more TG. However, the specific mechanism should be verified in the future.

## Conclusion

The experimental results show that konjac or exercise can significantly reduce body and liver weight, improve lipid metabolism. Moreover, the combined intervention of konjac and aerobic exercise was better than konjac or exercise, respectively. The mechanism of lipid metabolism regulation by exercise and konjac may decrease the LXRα and SREBP-1c protein expression levels to increase the ApoA5 expression and metabolize more TG (Fig 9). However, the specific mechanism should be verified further. Our findings demonstrate that konjac and exercise can improve lipid metabolism and oxidative status in obesity rat’s model induced by a high-fat diet.

**Fig 9.**
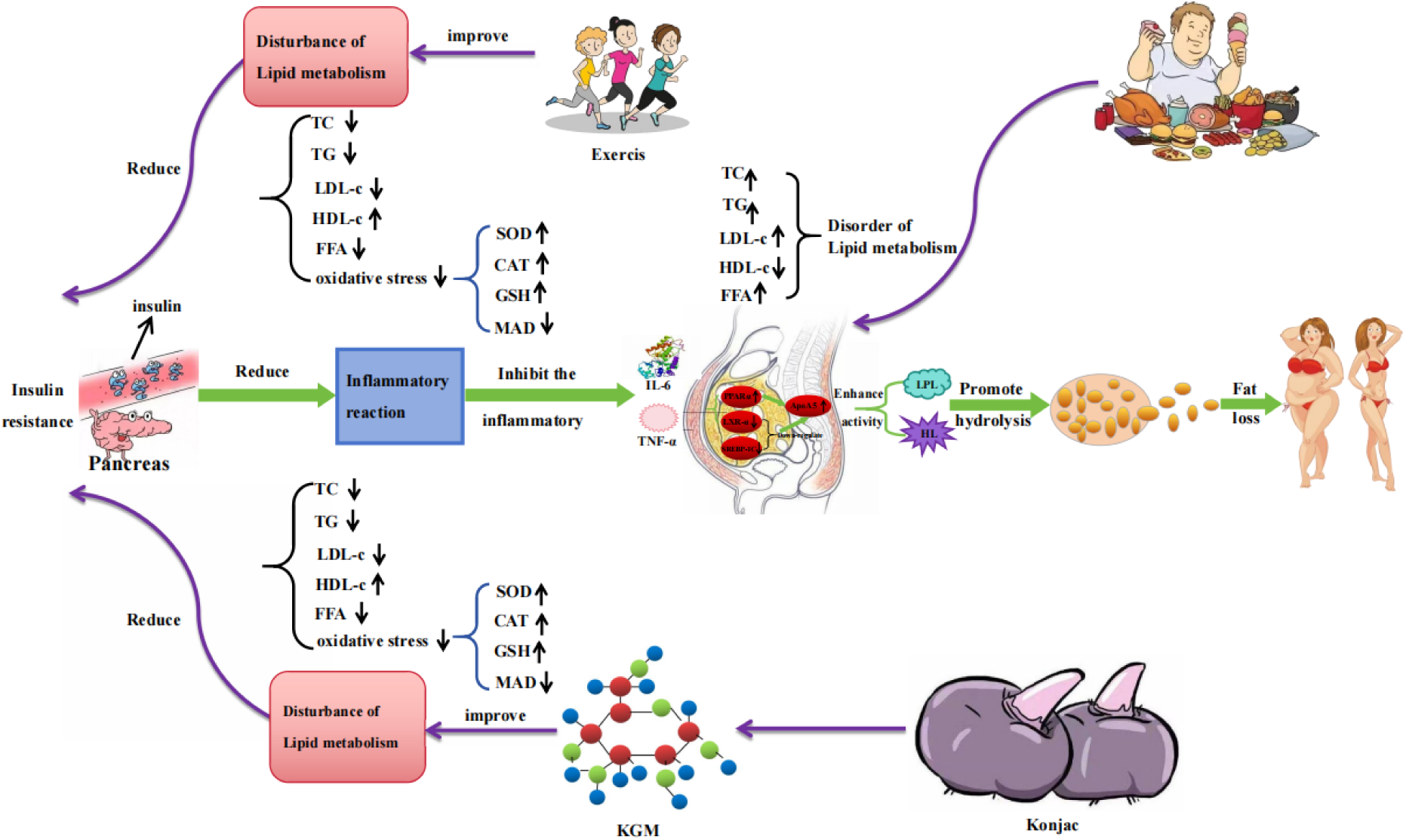
Effects and Mechanisms of Exercise and Konjac on Lipid Metabolism in Male Rats.

## Funding information

This study was funded by the National Natural Science Foundation of China (grant numbers 31500961, 61575065, 31771256), Graduate Student Innovation and Practical Ability Training Program of Xi ‘an Shiyou University (grant numbers: YCS23214339,YCS23241019), and innovation and Entrepreneurship Training Program for College Students of Xi’an Shiyou University (grant numbers:202110705018).And this study was partially funded by the Graduate Innovation and Practical Ability Training Project of Xi’an Petroleum University (Approval No. YCS23214339) and the Regular Project of Shaanxi Provincial Sports Bureau (Approval No. 240398,240223).

## Author contributions

Ling Ruan: conceived research, performed experiment, analyzed data. Ling Ruan, Shoubang Li, Guanghua Wang and Zhenqing Lv: performed experiment, revised manuscript and funding. Kun You and Guanghua Wang: revised tables and figures. Menglin Chen, Siyuan Liu: performed experiment. Lu Pu, Yun Pang: proofread the full text. Xinyue Liu, Siyu Liu: draw diagram.

## Conflict of Interest

“Competing interests: The authors declare there are no competing interests.”

## Data Availability Statement

The data presented in this study are available on request from the corresponding author. The data are not publicly available due to ethical restrictions.

## Acknowledgments

We thank all the staff for their efforts in this study. We are also grateful to the laboratory members for technical support during the experiment.

## Notes

### Competing Interest Statement

The authors have declared no competing interest.

